# Label Noise in Pathological Segmentation is Overlooked, Leading to Potential Overestimation of Artificial Intelligence Models

**DOI:** 10.1101/2025.02.18.638843

**Authors:** Kenji Harada, Yuichiro Nomura, Daisuke Komura, Shumpei Ishikawa, Shingo Sakashita

## Abstract

Artificial intelligence (AI) has transformed medical imaging, driving advancements in radiology and endoscopy. Semantic segmentation, a pixel-level technique crucial for delineating pathological features, has become a cornerstone of digital pathology. Pathology segmentation AI models are often trained using annotations generated by pathologists. Despite the meticulous care typically exercised, these annotations frequently contain empirical label noise. However, the specific types of label noise in pathology data and their impact on AI model training remain inadequately explored. This study systematically investigated the effects of label noise on the performance of pathology segmentation models. Using publicly available datasets and a breast cancer semantic segmentation dataset, modules were developed to simulate four types of artificial label noise at varying intensity levels. These datasets were used to train deep learning models with encoder-decoder architectures, and their performance was evaluated using metrics such as the Dice coefficient, precision, recall, and intersection over union. The results indicated that models were highly susceptible to overfitting label noise, particularly boundary-dependent noise such as dilation and shrinkage. Discrepancies were identified between apparent performance scores obtained under real-world conditions and true performance scores derived using clean test data. This overestimation risk was most pronounced for datasets containing boundary-altering noise. Furthermore, random combinations of noise types and levels significantly impaired model generalization. This study underscores the critical importance of addressing label noise in pathology datasets. It is proposed that future efforts focus on developing standardized methods for quantifying and mitigating label noise, along with creating robust benchmarks using noise-inclusive datasets. Enhancing annotation quality and addressing label noise can improve the reliability and generalizability of AI in pathology, facilitating broader clinical adoption.

## Introduction

The advent of artificial intelligence (AI) in healthcare has significantly transformed numerous medical fields. Recent advancements in radiology and endoscopy have notably enabled the integration of AI-based applications into clinical practice.^1^ In pathology, AI development for both clinical and research purposes is rapidly gaining attention.^2–4^ Among the critical AI techniques employed in pathology is semantic segmentation, a pixel-level method widely utilized to delineate lesions and other pathological features in medical images.^5,6^ Semantic segmentation is pivotal in digital pathology, as it facilitates the precise identification of tissue structures, cellular components, and pathological features in histological images. Detailed segmentation, such as detecting microscopic tumor cells or identifying vascular invasion, is essential for tasks where errors are intolerable. Recent progress in phenotypic and functional research on cellular and tissue structures, including analyses of protein expression, distribution, and complex tumor characteristics, depends on generating precise cell-level segmentation data.^7^

The training of pathology segmentation AI models often relies on annotations made by pathologists and data derived from immunohistochemical staining—both requiring considerable effort to produce.^8–13^ However, despite the meticulous care taken during the annotation process, these annotations frequently contain variability and errors, commonly referred to as label noise.^14^ Label noise in pathology arises from several sources. First, the inherent variability in human expert annotations contributes significantly to this noise.^15^ Even with common expertise, different pathologists may produce varying segmentations for the same image due to subjective interpretation, fatigue, or differing levels of experience. Second, the complexity of histological images—characterized by heterogeneous textures, overlapping structures, and subtle morphological changes—can create ambiguous regions that are difficult to annotate consistently. Additionally, manual annotation is time-consuming and labor-intensive, often leading to rushed or incomplete annotations that exacerbate label noise. Dilation and omission noise, referring to regions extending beyond the actual ground truth or false-negative areas where lesions are not annotated, respectively, are frequently encountered in this context.

The impact of label noise on the performance of segmentation models has been recognized across various domains. For example, Rahaman et al. investigated the effects of artificial label noise on AI models designed to identify water areas in satellite images, exploring noise types such as Gaussian, translation/shift, rotation, and mirroring/flipping.^16^ Similarly, studies using magnetic resonance images have examined the influence of label noise types, including random warp, constant shift, and random crop.^17^ These studies demonstrated that label noise significantly degrades model performance. Additionally, Luo et al. reported that dilation and erosion in fluorescence microscopy binary mask images reduce model segmentation performance.^18^ However, these studies lacked detailed analyses of the effects of different noise types and intensities or comparative evaluations of model generalizability. Moreover, limited validation of pathological segmentation data has been conducted, and the specific impact of label noise on pathological AI models remains unclear. Therefore, in addition to investigating label noise in public pathology data, this study aimed to elucidate its impact on the training of pathology segmentation AI and its effect on generalizability. Specifically, various levels and types of artificial label noise were introduced into a dataset initially considered to have minimal label noise, and the resultant effects were evaluated by comparing actual performance outcomes.

## Materials and Methods

### Datasets

We used four publicly available datasets to evaluate the presence of label noise: WSSS4LUAD,^9^ HuBMAP,^10^ CAMELYON,^11^ and GlaS^12^ (Fig. 1). The Breast Cancer Semantic Segmentation (BCSS) dataset was used to analyze artificial label noise.^13^ This dataset consists of 151 histopathological images of breast cancer, 0.25 microns per pixel, annotated for 21 different tissue components that include tumor regions, stroma, and inflammatory cells. This dataset was selected specifically for its detailed annotations, which allowed us to introduce controlled levels and types of artificial label noise. Labels with “angioinvasion (GT_code = 19)” and “dcis (GT_code = 20)” were combined as “tumor (GT_code = 1)”, because of the presence of tumor cells in both regions. The lesion labels were then binarized for model training and subsequent experiments. All datasets were reviewed by KH, and SS reviewed a representative example of the dataset.

**Figure 1.**
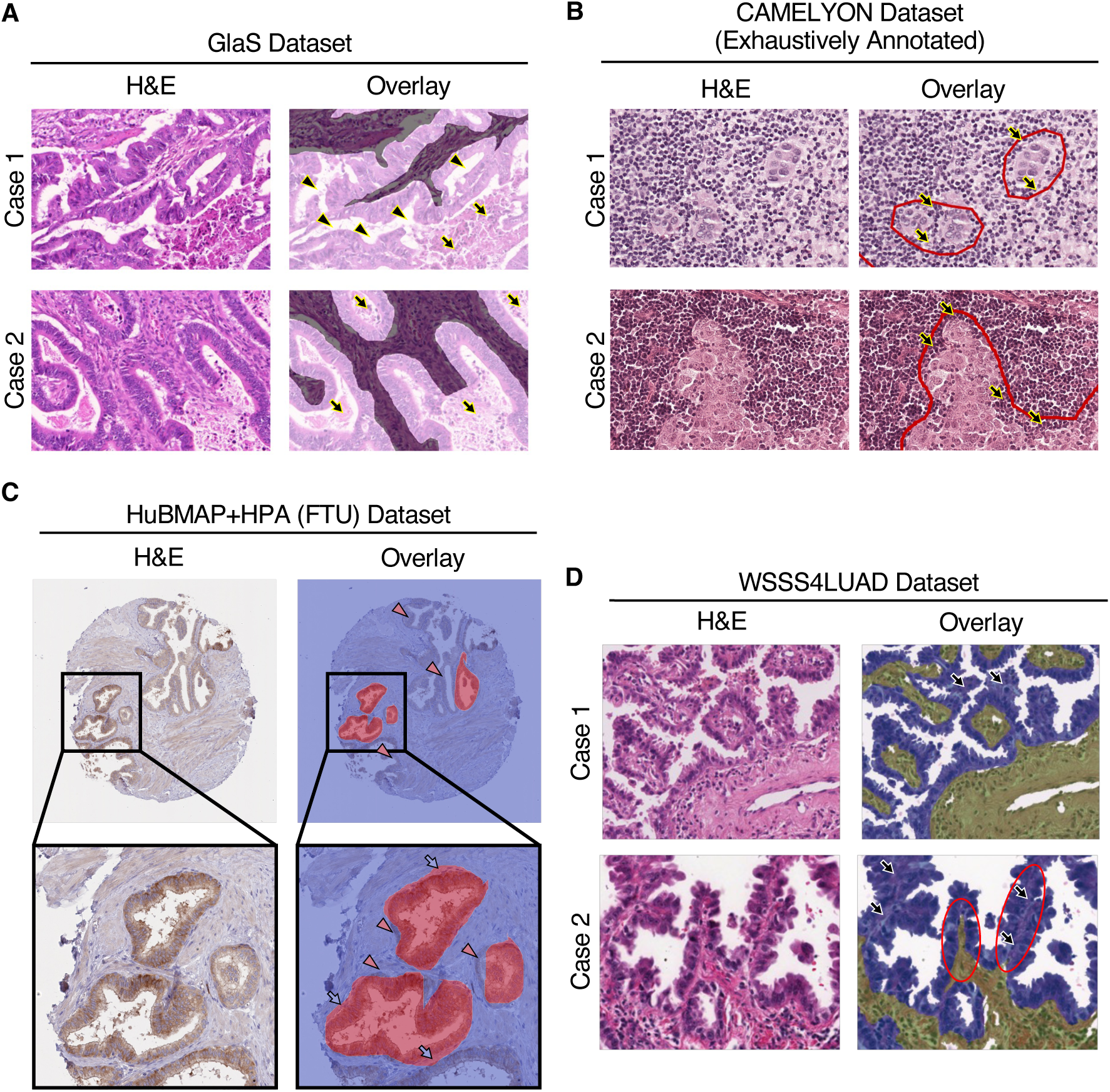
Public pathological segmentation data containing a variety of label noise. (A) Representative data from the GlaS dataset.^12^ The white region indicates the area labeled positive. Lumen (arrow head) and necrotic tissue areas (arrow) observed within the annotations. (B) Representative data of exhaustively annotated CAMELYON dataset.^11^ The red circle indicates the provided ground truth for the tumor cells. Non-tumor cells, such as lymphocytes (arrow) were also labeled in the tumor regions. (C) Representative data from the HuBMAP + HPA dataset.^10^ The red area indicates the region labeled positive. Unannotated structures (arrow head; both omission and shrinkage) and dilated annotation boundaries (arrow) were identified. (D) Representative data from the WSSS4LUAD dataset.^9^ The yellow and blue indicate provided annotations of the stroma and tumor areas, respectively. Instances of false positive regions (arrow) for tumor cells and inconsistencies in positive region annotations within the same image (red circle) were observed.

### Artificial label noises

To assess the effects of different types of label noise on pathology segmentation, we implemented and applied four types of artificial label noise: dilation, shrinkage, omission, and additive. These noise types were designed to simulate common annotation errors in histopathological images, and were introduced into the BCSS dataset as follows:

Dilation Noise: This noise simulates the expansion of annotated regions beyond their true boundaries. The implementation involved dilating the binary mask using a specified number of iterations, followed by smoothing the contours to avoid sharp edges of the mask. The number of iterations was randomly selected to span between 10–30 for level 1, 30–50 for level 2, and 50–150 for level 3. The smoothing parameter epsilon values, used to smooth the boundaries of the dilated regions, were set between 10–20 for level 1, 20–30 for level 2, and 30–50 for level 3. The original state was restored outside the region of interest (GT_code = 0).

Shrink Noise: Shrink noise represents the reduction of annotated regions. This noise was implemented by iteratively shrinking the mask through binary erosion. In this process, the implementation ensured the avoidance of omission noise by retaining small areas below a specified threshold. This process was repeated for a specific number of iterations, based on the noise level. The number of repeat processes was set to 1–5 for level 1, 5–9 for level 2, and 9–13 for level 3.

Omission Noise: Omission noise simulates the complete omission of certain annotated regions, which typically leads to false negatives. This was achieved by removing small regions that fell below certain area thresholds from the binary mask: 10,000–50,000, 50,000–100,000, and 100, 000–200,000 for levels 1, 2, and level 3, respectively.

Additive Noise: This type of noise introduces new incorrect regions into the annotation, typically leading to false positives. The implementation involved adding random geometric shapes (e.g., circles, ellipses, and polygons) of various sizes to regions that were originally negative. Specifically, 3–8 shapes were added for level 1 (each with an area of 20,000–60,000 pixels), 3–8 shapes for level 2 (each with areas of 60,000–120,000 pixels), and 5–8 shapes for level 3 (each between 120,000–160,000 pixels). The types of shapes (e.g., circles, ellipses, or polygons) were chosen randomly. The locations and sizes of these shapes were controlled to ensure that they did not overlap with true-positive regions, thereby avoiding dilation contamination.

The parameters for each noise type (e.g., iteration count and epsilon for dilation, number of repeats for shrinkage, area thresholds for omission, and number and size of regions for additive noise) were randomly generated based on predefined ranges corresponding to different noise levels (1, 2, and 3). This module is available on GitHub (at: https://github.com/kenjhara/noisypatho).

### BCSS dataset preparation and augmentation

The original histopathological images in the BCSS were divided into smaller patches of 512 × 512 pixels in size. A total of 6207 individual patch images were obtained from 151 total cases, The dataset was randomly split into three subsets: 80% of the images were used for training, 10% were used for validation, and the remaining 10% were reserved for testing. The dataset was pre-processed to normalize the image sizes and apply the appropriate augmentations. For the training phase, we implemented data augmentation techniques using the Albumentations library (version 1.3.1), including horizontal and vertical flips (each with a 50% probability), shift/scale/rotate transformations (with shift_limit = [–0.1, 0.1], scale_limit = [–0.1, 0.1], rotate_limit = [–90°, 90°], p = 1), and random brightness/contrast adjustments (brightness_limit = brightness_limit = [–0.1, 0.1], contrast_limit = [–0.1, 0.1], p = 0.5). These augmentations were applied to increase the diversity of the training data and enhance the model’s robustness. No augmentation was applied during the validation and testing phases, to ensure that this was performed on unaltered images.

### Model architecture

The training model was a neural network with an encoder-decoder structure. The encoder backbone consisted of pre-trained convolutional neural networks such as Resnet50^19^ or EfficientNet-b4.^20^ The decoder models included U-Net,^21^ U-Net++,^22^ FPN,^23^ PSPNet,^24^ and DeepLabV3+.^25^ The number of output classes was set to 2 (background and lesion). The final layer used a sigmoid activation function to perform the binary segmentation tasks.

### Training and validation

The models were trained using a combination of the dice loss and intersection over union (IoU) metrics as the primary evaluation criteria. Dice loss was selected for its effectiveness in terms of handling class imbalances, which are common in medical image-segmentation tasks. IoU was used to measure the overlap between predicted and true segmentation masks. We used the Adam optimizer, with an initial learning rate of 1 × 10^−3^ that was later decreased by a factor of 10 after a specified number of epochs. The training was monitored via early stopping, with a patience of 10 epochs. The training process was conducted for a maximum of 100 epochs, and the best-performing model (based on IoU validation) was saved for each configuration.

### Model evaluation

The models were evaluated using the testing dataset. The predicted segmentation masks were compared to the original ground truth or segmentation with artificial label noise using standard metrics that included precision, recall, dice coefficient, and IoU.

### Implementation

The artificial label noises were implemented using Python version 3.10.12, scipy version 1.10.1, pillow version 9.2.0, and opencv-python version 4.6.0.66. The neural networks were trained using PyTorch 2.0.1+cu117. The analysis and model training in this study were executed on Ubuntu 20.04 Linux system with an A100 GPU (NVIDIA, Santa Clara, CA, USA) and Ubuntu 18.04.5 Linux system with a GeForce GTX 1080 Ti GPU (NVIDIA).

## Results

### Public pathological segmentation data contain a variety of label noise

A number of pathological datasets with segmented data are currently available publicly, each created with different research objectives that have led to variations in the target cells or tissues. We therefore conducted a comprehensive manual examination of the annotations to assess the presence and extent of label noise present within them. Because it was difficult to quantitatively evaluate the noise present in the ground truth, a visual evaluation was performed by the authors. The GlaS dataset, which provides colorectal epithelial segmentation annotated by pathologists, was found to contain lumen and necrotic tissues within the annotations (Fig. 1A).^12^ The CAMELYON dataset, which includes detailed tumor cell annotations for lymph node metastases in breast cancer cases, divides the data into exhaustively annotated and non-exhaustively annotated categories based on annotation quality.^11^ However, even within the exhaustively annotated cases, non-tumor cells such as lymphocytes were observed to be incorrectly included in the positive regions (Fig. 1B). The HuBMAP + HPA dataset, used in the “HuBMAP + HPA— Hacking the Human Body” competition on Kaggle, is a large-scale dataset that includes segmentation at the cellular level for five organs: kidney, prostate, colon, spleen, and lung.^10^ Label noise was evident across the dataset, with issues such as unannotated structures, lumen contamination, and inconsistent annotation boundaries being identified (Fig. 1C). The WSSS4LUAD dataset, focused on lung adenocarcinoma, includes tissue semantic segmentation data.^9^ Upon closer inspection, instances of false-positive regions for tumor cells and inconsistencies in positive region annotations within the same image were observed (Fig. 1D). These findings indicated that current publicly available datasets exhibit various types of label noise—including dilation, shrinkage, and omission—with varying degrees of noise present between the datasets and individual cases. Although the creation of segmentation data for pathology is inherently labor-intensive and subject to certain limitations, the presence of label noise is a critical issue that must be carefully considered, as it can potentially impact AI training outcomes.

### Generation of artificial label noise

We generated artificial label noise to investigate its impact on AI performance. We developed modules to create three types of label noise—shrink, dilation, and omission — based on observations from publicly available datasets. In addition to these noises, we applied additive noise, which is the counterpart to omission noise. These four types of noises were added to the annotated BCSS dataset, which we determined to have one of the lowest levels of label noise among the publicly available datasets we investigated (Fig. 2A; Supplementary Fig. S1A,B). Each type of label noise was introduced at three different levels, allowing for a range of noise intensities (Fig. 2B,C; Supplementary Fig. S1C,D). In addition, to simulating conditions that more closely mimicked real-world scenarios, we created a combined pattern of dilation and omission noise, which is commonly observed in pathological data. When comparing the datasets with and without artificial label noise, distinct patterns in precision, recall, dice coefficient, IoU, and the ratio of noise area to non-tumor (i.e., class 0) areas were observed for each type of label noise (Fig. 2D; Supplementary Fig. S2). In total, we successfully generated 15 dataset variations, encompassing five types of label noise across three levels of intensity.

**Figure 2.**
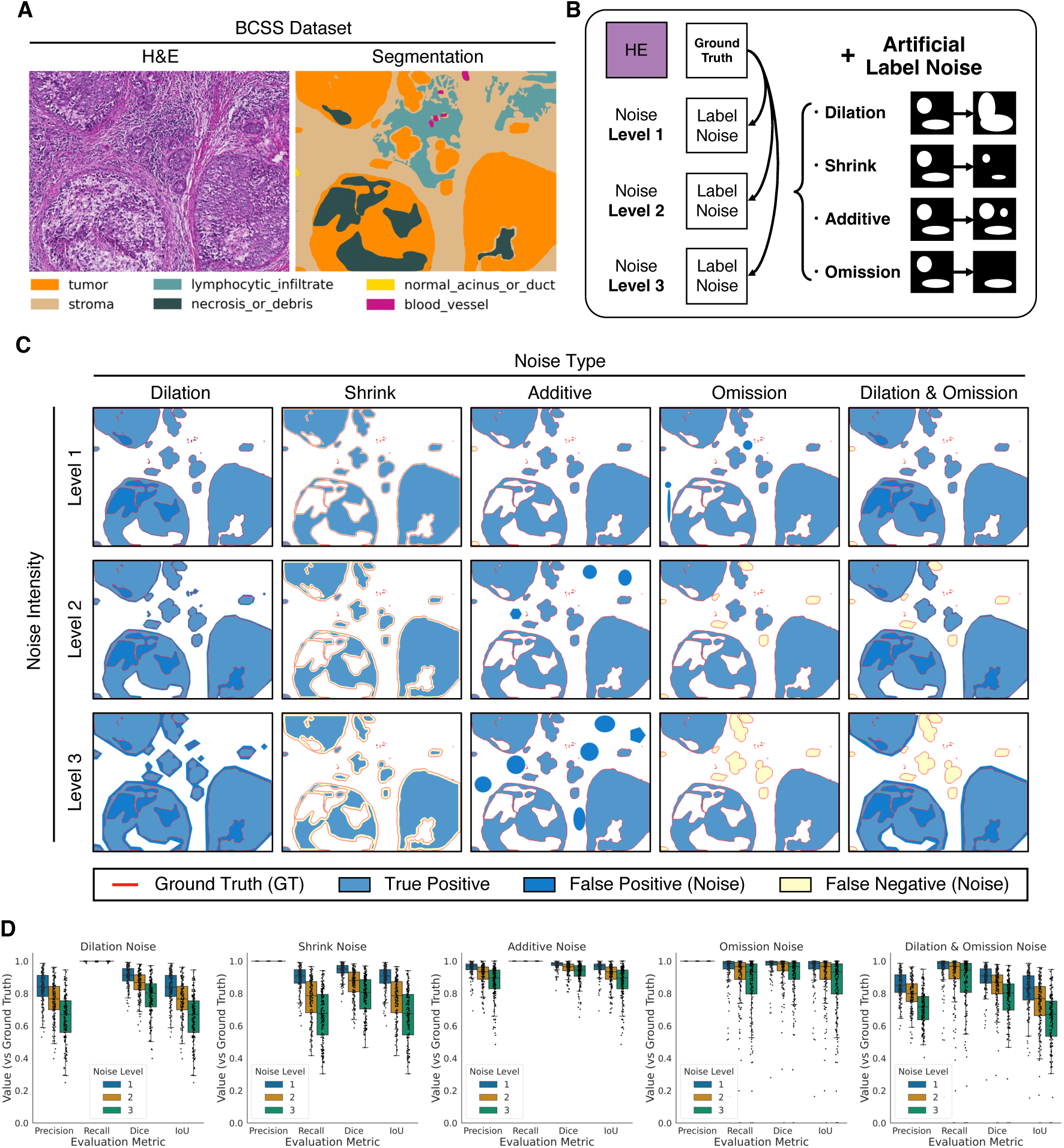
Artificial label noise. (A) A representative image from the BCSS dataset.^13^ (B) An overview of the artificial label noise module. (C) Representative mask images showing the ground truth boundary (red line) alongside the noisy annotations generated by our module. Regions where the noisy annotation matches the ground truth are displayed in light blue. Areas where the ground truth is missing in the noisy annotation (i.e., under-segmentation) are shown in yellow. Areas where the noisy annotation includes regions not present in the ground truth (i.e., over-segmentation) are shown in dark blue. (D) The precision, recall, dice, and intersection over union (IoU) scores were categorized by different types and levels of label noise vs the ground truth. The box represents the quartiles of the dataset, while the whiskers extend to display the rest of the distribution. Each dot represents an individual case.

### Label noise has a significant impact on model training

To assess the effects of label noise on AI model performance, we conducted training on datasets with clean labels and 15 artificially generated noisy datasets. We trained 10 models on each dataset, using combinations of two encoder architectures and five decoder types (Fig. 3A). The performance scores of the models trained on clean labels is detailed in Supplementary Table S1. We found that the combination of EfficientNet-b4 with either U-Net or DeepLabV3+ consistently yielded the highest accuracy. These two models demonstrated high predictive accuracy for the testing dataset when trained on clean labels (Fig. 3B; Supplementary Fig. S3A,B). When the models were trained using datasets with various types of introduced label noises, the predictions of the models generally reflected the same types of noise that were present in the training data (Fig. 3C); however, this was not the case for additive noise (Supplementary Fig. S4). Additionally, a clear drop in model performance was observed as the severity of the label noise increased (Fig. 3D). These results indicate that label noise is likely to have a significant effect on model learning.

**Figure 3.**
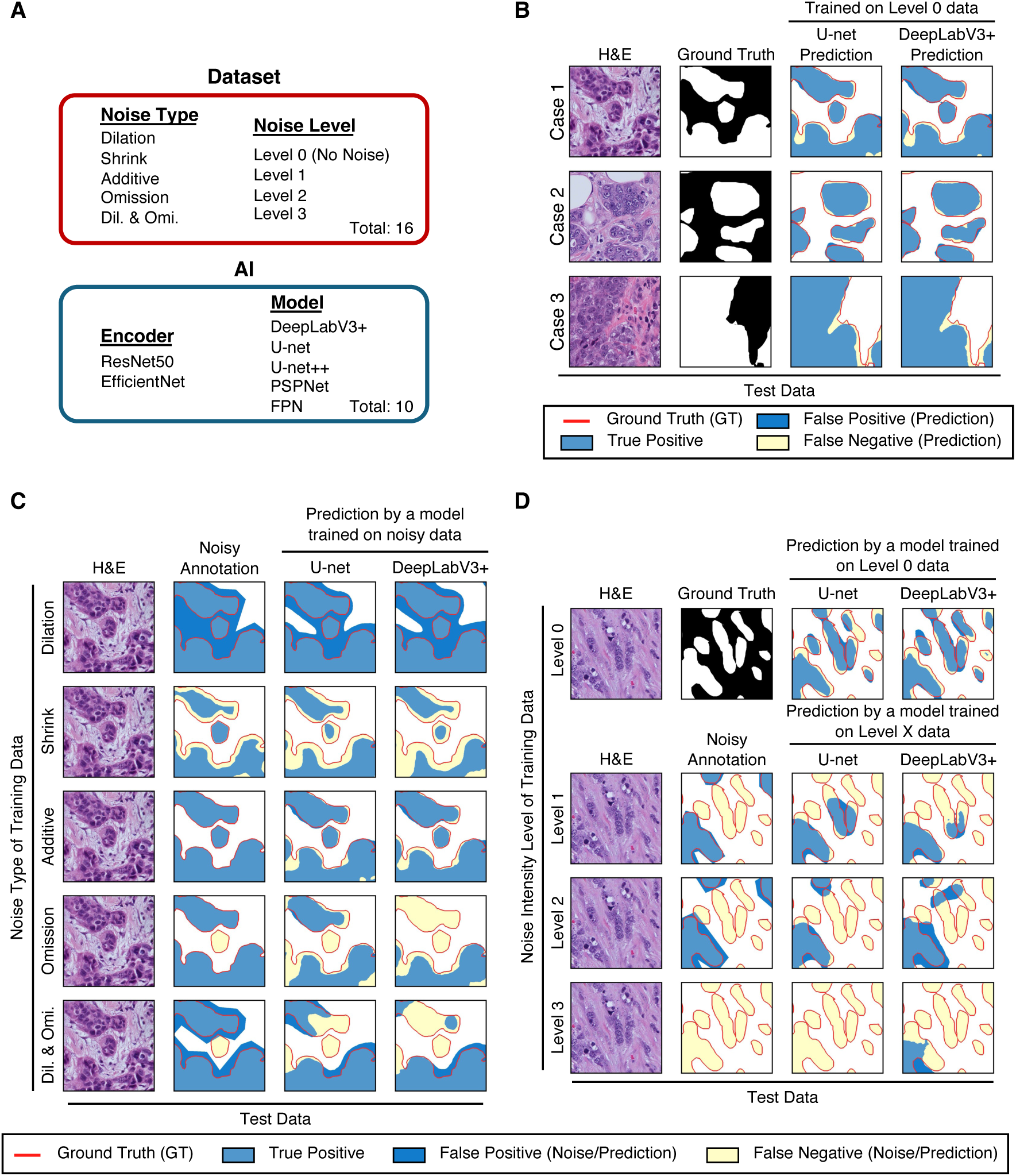
Visualization of the impact of label noise on model training. (A) Overview of the training. A total of 160 combinations were generated, using 16 datasets across 10 different models. (B) Visualization of the predictions made by the models trained on the level 0 dataset (no noise) for the test dataset. The white areas represent tumor regions in the ground truth mask images, while the black ones indicate non-tumor regions. The ground truth boundary is marked with a red line in the prediction mask images. Regions where the model’s predictions align with the ground truth are shown in light blue. Areas where the model fails to identify the ground truth (i.e., under-segmentation) are represented in yellow, while regions where the model includes predictions that were not found in the ground truth (i.e., over-segmentation) are displayed in dark blue. (C) Visualization of the predicted results on the test data using models trained with level 2 noisy labels, categorized by type of noise. For the noisy mask images, the ground truth boundary is marked with a red line. Regions where the noisy annotations align with the ground truth are shown in light blue. Areas where the ground truth is missing in the noisy annotations (i.e., under-segmentation) are represented in yellow, and regions where the noisy annotations include areas that were not found in the ground truth (i.e., over-segmentation) are displayed in dark blue. The coloring pattern of the predictions are the same as in (B). (D) Visualization of the prediction results on the testing dataset using models trained with either the level 0 dataset (no noise), or one with dilation and omission label noise, categorized by each level of noise. The coloring patterns are the same as in (B,C). Dil., Dilation, Omi., Omission.

### Label noise often goes underestimated in pathology segmentation

We evaluated model performance on the testing dataset using two approaches. The first involved testing data that contained noise at comparable levels to the training data, allowing us to calculate apparent real-world performance scores (Fig. 4A; Supplementary Fig. S5A). The second approach used clean labels for the testing data, in order to provide more true and idealized performance scores (Fig. 4B; Supplementary Fig. S5B). Our comparison revealed that, for models trained on datasets with dilation, shrinkage, or combined noise with both dilation and omission, the apparent scores were often inflated relative to the true ones—indicating an overestimation of model performance (Fig. 4C,D; Supplementary Fig. S5C,D). By contrast, models trained with additive noise showed higher true performance scores vs apparent scores. For omission noise, there was little difference between the two scoring methods. These results indicate that such models may overestimate identified targets if the dataset includes noise that alters the object’s boundaries, such as dilation or shrinkage.

**Figure 4.**
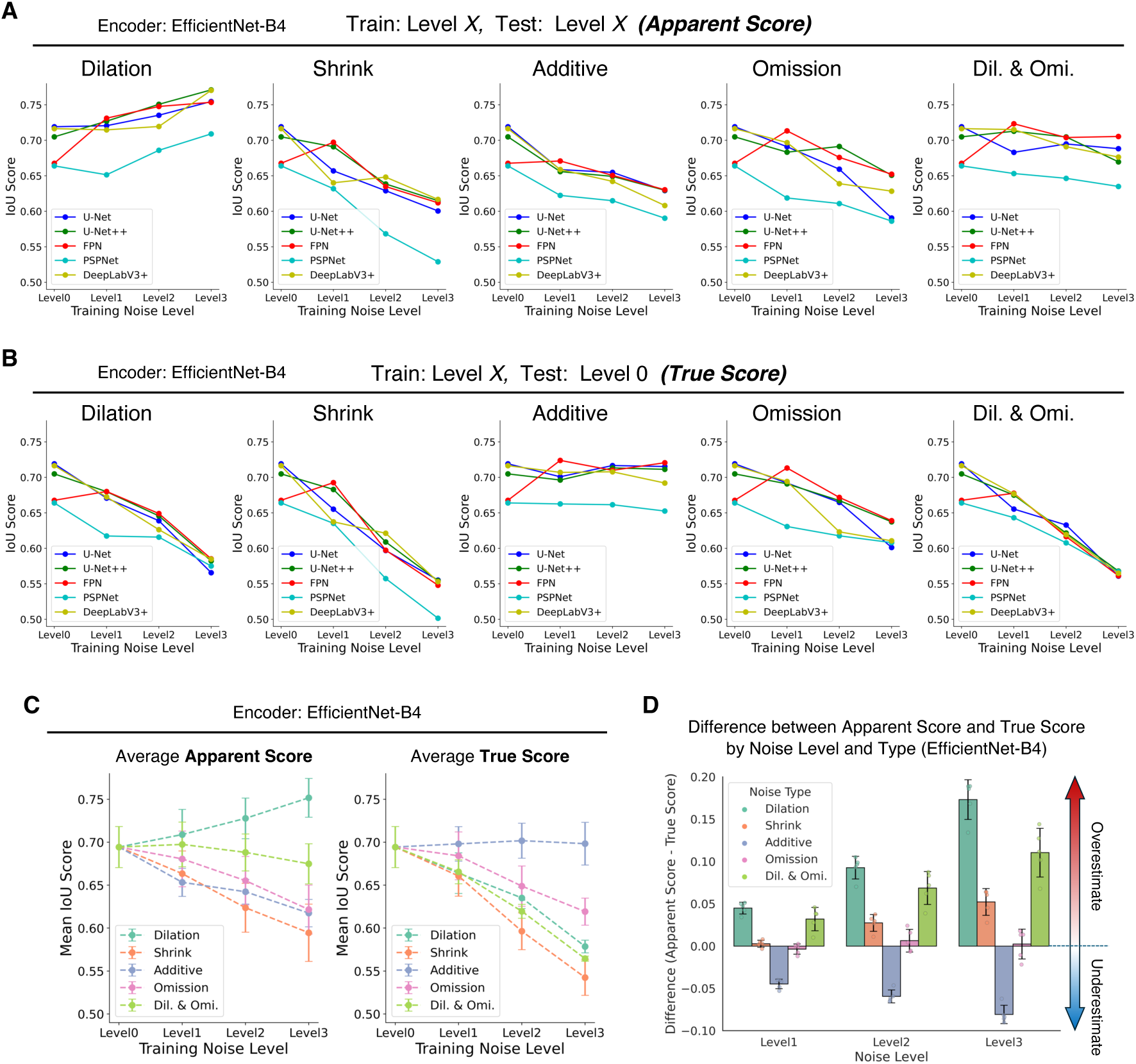
Label noise often goes underestimated in pathology segmentation. (A) Apparent scores were calculated using the testing dataset for all models with EfficientNet-b4, showing the noise level used for the training on the x-axis and the IoU score on the y-axis, summarized by noise type. (B) True scores were calculated using the test dataset for all models with EfficientNet-b4, showing the noise level used for the training on the x-axis and the IoU score on the y-axis, summarized by noise type. (C) Average apparent and true scores for all models shown in (A) and (B), showing the noise level used for the training on the x-axis and IoU score on the y-axis. Points and error bars represent mean ± standard deviation values. (D) Difference between apparent and true scores by noise level and type. Each dot represents an individual model trained, with the level of noise on the x-axis. Error bars represent mean ± standard deviation values. Positive values indicate overestimation, 0 indicates evaluation match, and negative values indicate underestimation. Dil., Dilation, Omi., Omission.

### Random label noise also impairs model training

In real-world scenarios, label noise is rarely uniform; rather, various types of noise are randomly distributed across the dataset. To simulate this, we generated datasets containing random combinations of label noise types and evaluated their impacts on model performance (Fig. 5A). Similarly to the results we observed when using uniform noise, model performance was found to decline as the intensity of random noise increased in the data (Fig. 5B). Furthermore, a significant gap was observed between the apparent and true scores, indicating that models trained on datasets containing random label noise may also be prone to overestimation (Fig. 5C). We observed a substantial drop in performance in another experiment wherein the datasets contained label noise of varying intensity levels, confirming that the presence of random noise severely affected model training (Fig. 5D–F).

**Figure 5.**
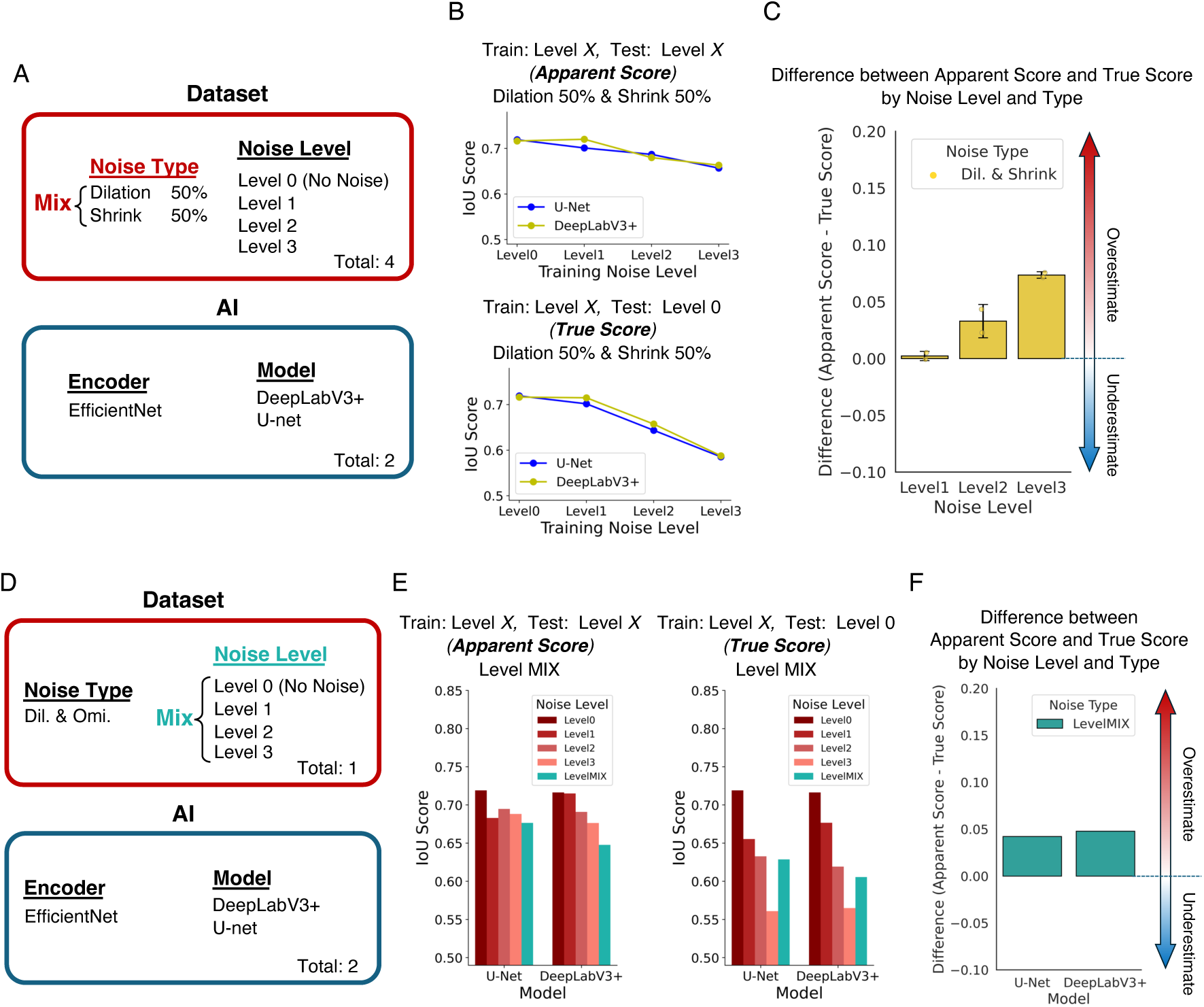
Random label noise also impairs model training. (A) An overview of the training using mixed random label noise of dilation and shrink. Eight combinations were generated, using four datasets across two different models. (B) Apparent and true scores were calculated using the test dataset for all models trained as shown in (A), showing the noise level used for the training on the x-axis and the IoU score on the y-axis. (C) Difference between apparent and true scores by noise level on models trained as shown in (A). Each dot represents an individual trained model, with the noise level on the x-axis. Error bars represent mean ± standard deviation values. Positive values indicate overestimation, 0 indicates evaluation match, and negative values indicate underestimation. (D) Overview of the training using mixed random label noise, by level of noise. Two combinations were generated, using single datasets across two different models. (E) Apparent and true scores were calculated using the test dataset for all models trained as shown in (D), with the models and their noise levels shown on the x-axis and IoU scores on the y-axis. (F) Differences between apparent and true scores on the models trained as shown in (D). Positive values indicate overestimation, 0 indicates evaluation match, and negative values indicate underestimation. Dil., Dilation, Omi., Omission.

## Discussion

This study investigated the presence of different types of label noise in publicly available pathology segmentation datasets, and demonstrated the significant impact of label noise on the training of AI models for pathology image segmentation. Specifically, random label noise caused by dilation, shrinkage, omission, or combinations thereof, was found to significantly decrease model performance. Furthermore, by comparing true scores with apparent ones, this study is the first to our knowledge to highlight the potential of label noise in terms of causing overestimations of model performance.

The label noise observed in existing datasets can be categorized into three primary types: dilation, shrinkage, and omission—each of which stems from factors such as the ambiguity of lesion boundaries and subjective biases in pathologist annotations. For instance, dilation may occur because of a tendency to mark boundaries more broadly for safety, whereas shrinkage may result from a cautious approach that focuses only on definitive regions. Omission, on the other hand, may stem from missed annotations of small or subtle lesions, potentially caused by fatigue, inattention, or lack of experience. Additive noise, which involves the erroneous labeling of non-lesion areas with visual characteristics as actual lesions, may also occur—although its classification as noise is still debated among experts in the field. In this study, we developed a module to generate artificial label noise artifacts that mimicked these specific noise types, which are prevalent in pathology but underexplored in other domains.^16,17,26,27^

Using 15 types of artificial label noise, we trained deep-learning models and compared their performances on our testing datasets. Figure 3C,D shows that the models were highly susceptible to overfitting label noise, which is consistent with the results of prior studies concerning label noise in image classification tasks.^28^ Notably, Han et al. demonstrated that deep learning models initially prioritize learning from clean labels but eventually memorize noisy labels as training progresses, in a phenomenon known as the “memorization effect”.

We then evaluated the IoU scores of the model predictions on the testing datasets using both clean and noisy labels, referred to as the true and apparent scores, respectively. Because it is generally unknown whether annotation data represent the expected ground truth, apparent scores are typically calculated under standard experimental conditions. True scores, however, are only calculable in experiments involving artificial label noise. A comparison of the true scores revealed that the models maintained their performance more optimally when exposed to noise types in the following order: additive, omission, dilation, and shrink. Additive noise, characterized by randomly introducing small, irregular class 1 regions into class 0 areas, likely avoided overfitting because the remaining class 0 regions were substantially larger than the introduced class 1 regions (Supplementary Fig. S2). Another possible explanation for this may be related to label noise location. In our experiments, additive noise artifacts were randomly generated in non-tumor regions, indicating that the characteristics of the target region were inconsistent. However, it can be assumed that actual additive noise is most likely generated in regions that are similar to the targets. If this is the case, it may have substantial impacts on the learning process. By contrast, our models that were trained with shrinkage and dilation noise exhibited lower performance, likely owing to the added noise in the indistinct mask boundary areas, which also exacerbated issues of class imbalances. These findings demonstrate the varied characteristics of different types of label noise, as well as their distinct effects on model performance.

Moreover, the differences between the apparent and true scores varied with the noise type and intensity, underscoring the potential for overestimation of model performance when label noise is present. This discrepancy is particularly pronounced for boundary-dependent noises such as dilation, shrinkage, and random variations. Our findings also corroborate certain existing challenges that have been identified when generalizing AI models, wherein models trained on specific datasets often perform poorly when given new data.^29,30^ This lack of generalizability can be attributed to the limited diversity and external validation of datasets. Many studies rely on only one or two data sources, which further exacerbates this issue. Although recent methods have been developed that facilitate the training of AI models using smaller datasets, manual annotation, and corrections remain essential—indicating that the challenges related to label noise have not yet been fully addressed. Our present results highlight the potential for overestimation when validation is restricted to limited data sources.

Pathology-specific label noise presents unique challenges that are not typically encountered in other imaging domains. For instance, slides stained with Hematoxylin and Eosin often exhibit ambiguous cellular boundaries, which can amplify the inherent variability in pathologist annotations and hinder the construction of large, high-quality datasets. Segmentation decisions can vary significantly even among experienced and well-trained experts, making it difficult to establish a single “correct” annotation.^31^ Thus, the definition of label noise in pathology remains inherently complex. Nevertheless, most pathologists can identify the subjective presence of label noise. Future efforts should focus on developing methods for both the qualitative and quantitative evaluation of label noise, particularly ones that mitigate boundary-dependent noise such as dilation and shrinkage. The development of standardized methods for not only quantifying but also correcting label noise, is therefore warranted in this field. This requires that annotated databases be created, firstly, that are designed to include various levels and types of label noise. As we demonstrated in this study, the introduction of artificial label noise partially facilitated this process. Furthermore, such databases would facilitate benchmarking for model robustness, paving the way for the development of more resilient AI systems. Constructing and evaluating large-scale databases that systematically record label noise will also provide researchers in the field with valuable resources. Ultimately, improving annotation quality and minimizing the effects of label noise are essential for fully realizing the potential of AI in the context of pathological image analysis.

This study systematically characterized the various types of label noise present in pathology datasets, and investigated their impact on model performance by simulating this type of noise. Our results demonstrated the significant influence of boundary-altering noise on AI model performance in this context. By comparing apparent scores to true ones, we highlighted the risk that the performance metrics of existing models may be overestimated. This study highlights the importance of recognizing and addressing label noise as a critical challenge during the development of pathological AI systems.

## Author Contributions

K.H. conceptualized the study; K.H. and Y.N. developed the methodology; K.H. performed the formal analysis; D.K., S.I., and S.S. provided the resources; K.H. and Y.N. wrote and prepared the original draft; D.K., S.I., and S.S. reviewed, and edited the manuscript; and K.H. and S.S. supervised the study and provided funding acquisition. All of the authors have read and approved the final submitted version of the manuscript.

## Acknowledgments

We thank Professor Norio Sakai (Department of Molecular and Pharmacological Neuroscience, Graduate School of Biomedical and Health Sciences, Hiroshima University) for his kind support and for allowing us to use his facilities. During the preparation of this work, the authors used ChatGPT to correct misspellings, grammar, and sentence structure to improve the English writing. After using this tool, the authors reviewed and edited the content as needed, and take full responsibility for the content of the publication. We would like to thank Editage (www.editage.jp) for English language editing.

## Data Availability

We used publicly available data from the WSSS4LUAD, HuBMAP, CAMELYON, GlaS, and BCSS databases. The modules developed in this study are available on GitHub (https://github.com/kenjhara/noisypatho). The remaining data supporting the findings of this study are available from the corresponding author upon reasonable request.

## Ethics Approval and Consent to Participate

This study did not involve the collection or use of any clinical specimens or patient data from our institution. All of the experiments were conducted exclusively using publicly available datasets. Ethical approval and informed consent were therefore not considered necessary for this study.

---

**Supplementary Figure S1.**
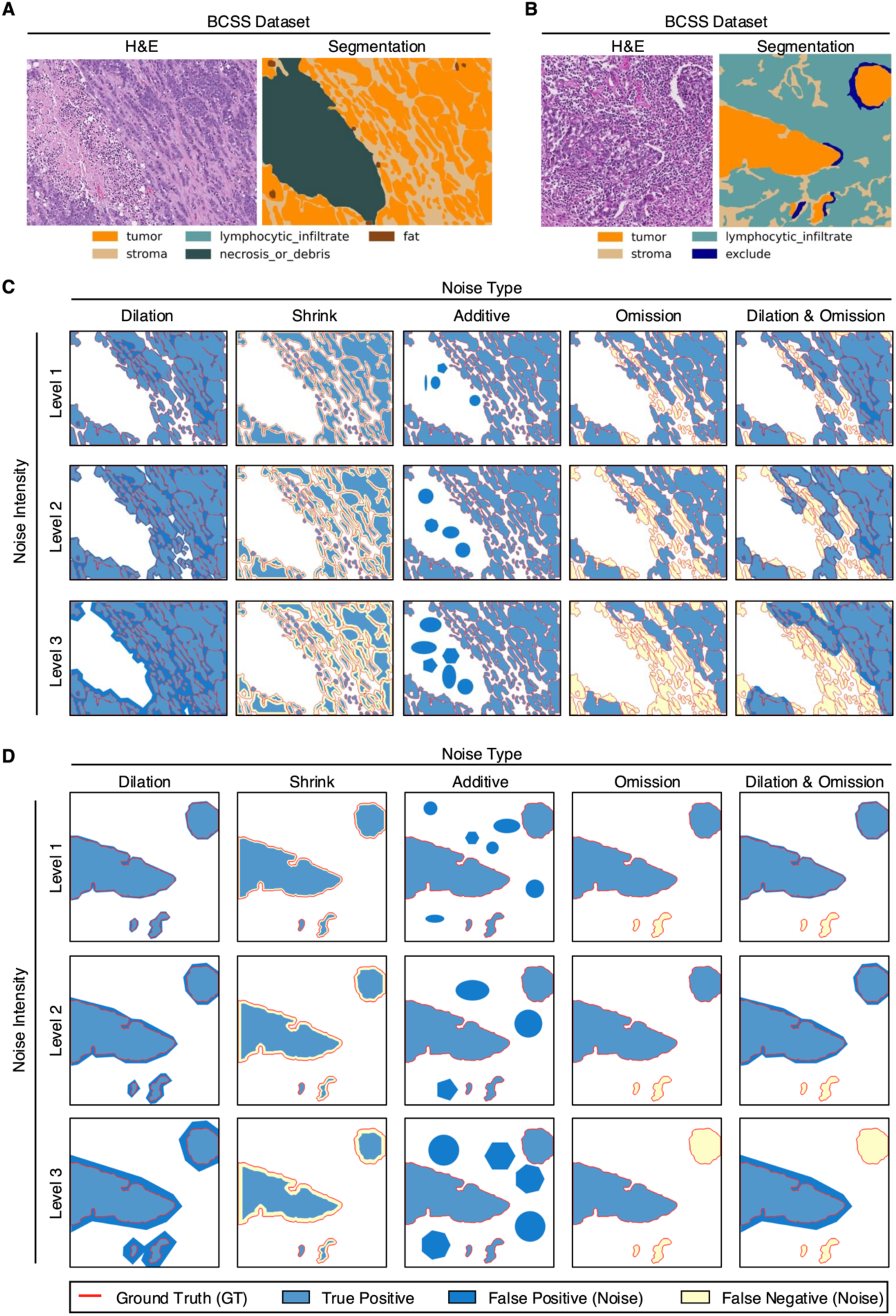
Artificial label noise. (A, B) Representative images from the BCSS dataset.^13^ (C, D) Representative mask images showing the ground truth boundary (red line) alongside the noisy annotations generated by our module. Regions where the noisy annotation matches the ground truth are displayed in light blue. Areas where the ground truth is missing in the noisy annotation (i.e., under-segmentation) are shown in yellow. Areas where the noisy annotation includes regions do not present in the ground truth (i.e., over-segmentation) are shown in dark blue.

**Supplementary Figure S2.**
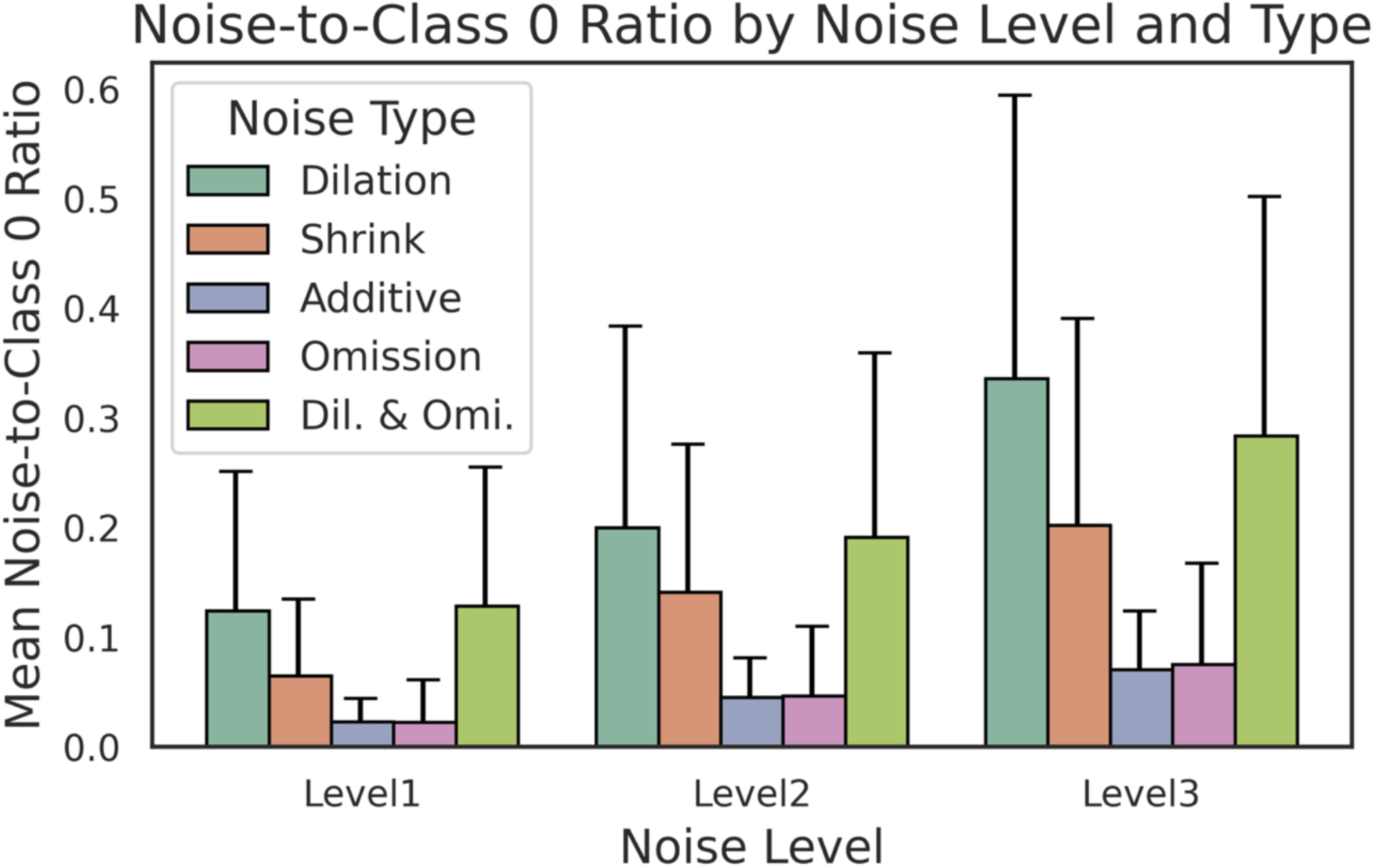
Ratio of Noise-to-Class 0 (non-tumor region). Ratio of the area increased or decreased by the introduction of artificial label noise (noise) to the area of non-tumor areas (class 0). Error bars are presented as mean + standard deviation.

**Supplementary Figure S3.**
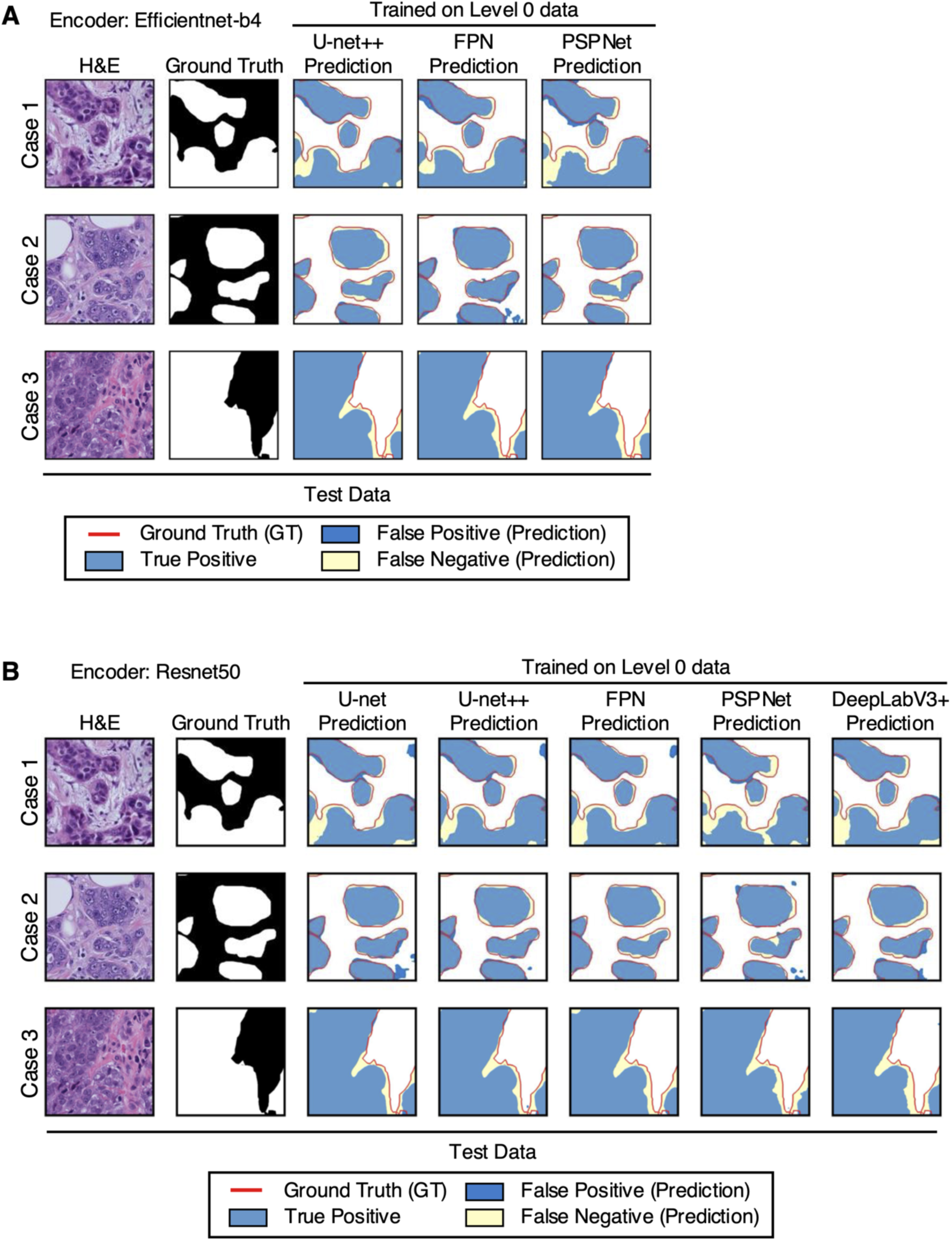
Visualization of predictions made by models trained on the level 0 dataset (no noise) for the test dataset. (A) Predictions made by the model using an EfficientNet-B4 encoder. (B) Predictions made by the model using a Resnet50 encoder. In the ground truth mask images, white areas represent tumor regions, while black areas indicate non-tumor regions. The ground truth boundary is marked with a red line in the prediction mask images. Regions where the model’s predictions align with the ground truth are shown in light blue. Areas where the model fails to capture the ground truth (i.e., under-segmentation) are represented in yellow, whereas regions where the model includes predictions absent from the ground truth (i.e., over-segmentation) are displayed in dark blue.

**Supplementary Figure S4.**
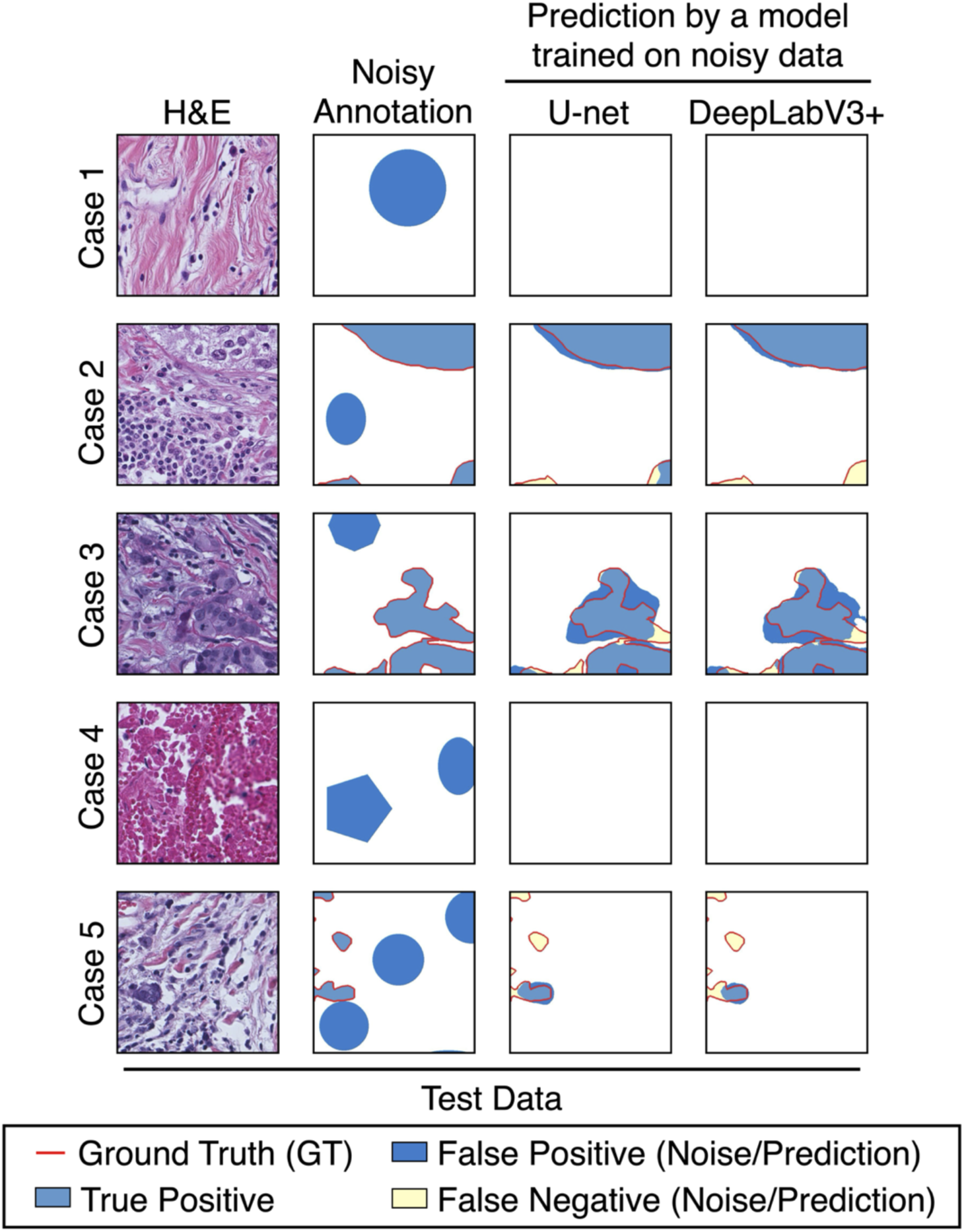
Visualization of the impact of additive label noise on model training. Visualization of the predicted results on the test data using models trained with level 1 additive label noises. For the noisy mask images, the ground truth boundary is marked with a red line. Regions where the noisy annotations align with the ground truth are shown in light blue. Regions where the noisy annotations include areas not found in the ground truth (i.e., over-segmentation) are displayed in dark blue. For the prediction mask images, the ground truth boundary is marked with a red line. Regions where the model’s predictions align with the ground truth are shown in light blue. Areas where the model fails to identify the ground truth (i.e., under-segmentation) are represented in yellow, and regions where the model includes predictions not found in the ground truth (i.e., over-segmentation) are displayed in dark blue.

**Supplementary Figure S5.**
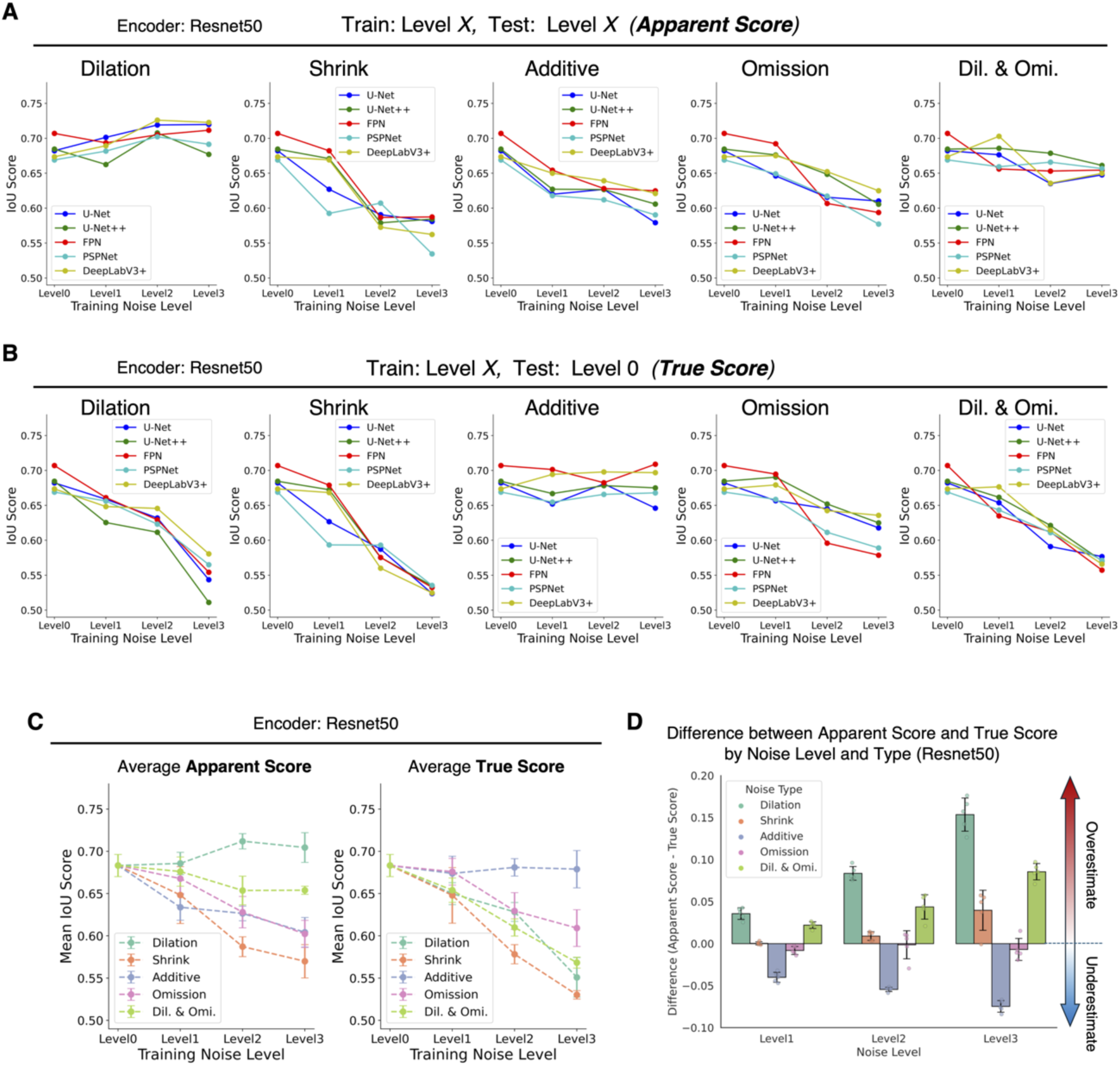
Data related to. Figure 4. (A) Apparent scores were calculated using the test dataset for all models with Resnet50, showing the noise level used for the training on the x-axis and the IoU score on the y-axis, summarized by noise type. (B) True scores were calculated using the test dataset for all models with Resnet50, showing the noise level used for the training on the x-axis and the IoU score on the y-axis, summarized by noise type. (C) Average apparent and true scores for all models shown in (A) and (B), showing the noise level used for the training on the x-axis and IoU score on the y-axis. Points and error bars represent mean ± standard deviation values. (D) Difference between apparent and true scores by noise level and type. Each dot represents an individual model trained, with the level of noise on the x-axis. Error bars represent mean ± standard deviation values. Positive values indicate overestimation, 0 indicates evaluation match, and negative values indicate underestimation. Dil., Dilation, Omi., Omission.

**Supplementary Table S1.** Performance of models trained on clean labels (Level 0 labels)

| Encoder | Model | Precision | Recall | Dice | IoU |
| --- | --- | --- | --- | --- | --- |
| Efficientnet-b4 | U-net | 0.875 | 0.803 | 0.784 | 0.719 |
|  | U-net++ | 0.855 | 0.804 | 0.770 | 0.705 |
|  | FPN | 0.814 | 0.791 | 0.737 | 0.668 |
|  | PSPNet | 0.805 | 0.783 | 0.735 | 0.664 |
|  | DeepLabV3+ | 0.867 | 0.802 | 0.780 | 0.716 |
| Resnet50 | U-net | 0.796 | 0.817 | 0.750 | 0.682 |
|  | U-net++ | 0.802 | 0.817 | 0.752 | 0.684 |
|  | FPN | 0.878 | 0.786 | 0.772 | 0.707 |
|  | PSPNet | 0.834 | 0.773 | 0.739 | 0.669 |
|  | DeepLabV3+ | 0.867 | 0.755 | 0.746 | 0.673 |

